## Supplemental Table 3 for "A complex of BRCA2 and PP2A-B56 is required for DNA repair by homologous recombination"

**Table S3.**

| ***BRCA2 in syringe/***  ***B56α in calorimetric cell*** | **ΔG**  **(*kcal mol^-1^*)** | **ΔH**  **(*kcal mol^-1^*)** | **-TΔS**  **(*kcal mol^-1^*)** | **K_D_**  **(*μM*)** | **n** |
| --- | --- | --- | --- | --- | --- |
| B56α/BRCA2*^1108-1126^* | -7.21 | -14.9 ± 0.20 | 7.69 | 5.17 ± 0.25 | 1.02 ± 0.01 |
| B56α/BRCA2*^1108-1126^* *L1114A, I1117A* | No binding | | | | |
| B56α/BRCA2*^1105-1129^* | -6.97 | -13.8 ± 0.25 | 6.87 | 7.76 ± 0.40 | 0.99 ± 0.01 |
| B56α/BRCA2*^1105-1129^* *pS1106* | -6.70 | -12.2 ± 0.30 | 5.50 | 12.3 ± 0.76 | 1.19 ± 0.01 |
| B56α/BRCA2*^1105-1129^* *T1116P* | -7.04 | -13.4 ± 0.32 | 6.35 | 6.92 ± 0.47 | 0.85 ± 0.01 |
| B56α/BRCA2*^1105-1129^* *D1106Q, D1123Q, D1128Q* | -7.40 | -15.4 ± 0.24 | 8.02 | 3.75 ± 0.22 | 0.86 ± 0.01 |
| B56α/BRCA2*^1105-1129^* *S1106R* | -7.55 | -14.4 ± 0.10 | 6.89 | 2.93 ± 0.09 | 0.95 ± 0.00 |
| B56α/BRCA2*^1105-1129^* *T1128I* | -7.00 | -16.2 ± 0.41 | 9.24 | 7.43 ± 0.43 | 1.01 ± 0.00 |
| B56α/BRCA2*^1113-1129^* | -7.39 | -18.0 ± 0.15 | 10.60 | 3.82 ± 0.12 | 1.01 ± 0.00 |
| B56α/BRCA2*^1113-1129^ pS1123* | -8.34 | -19.3 ± 0.09 | 11.00 | 0.77 ± 0.03 | 0.92 ± 0.00 |
| B56α/BRCA2*^1113-1129^ pT1128* | -7.99 | -17.7 ± 0.13 | 9.66 | 1.38 ± 0.04 | 0.94 ± 0.00 |
| B56α/BRCA2*^1113-1129^ pS1123, pT1128* | -8.62 | -19.3 ± 0.14 | 10.70 | 0.48 ± 0.03 | 0.91 ± 0.00 |
